# Identification of neuronal ensembles involved in remote fear memory extinction impairments

**DOI:** 10.1101/2023.09.17.558133

**Authors:** Kamil F. Tomaszewski, Kacper Łukasiewicz, Magdalena Robacha, Kasia Radwańska

**Affiliations:** Laboratory of Molecular Basis of Behavior, the Nencki Institute of Experimental Biology of Polish Academy of Sciences, Warsaw, Poland; Psychiatry Center, Medical University of Białystok, Poland; Department of Molecular and Systemic Neurophysiology, RWTH, Aachen, Germany

**Keywords:** PTSD - Posttraumatic stress disorder_1_, fear extinction_2_, remote fear extinction_3_, c-Fos_4_, CaMKII alpha_5_, remote memory_6_

## Abstract

Remote memory consolidation, extinction, and its impairments have been of interest to researchers for years, especially due to its clinical relevance in patients with emotional disorders, like PTSD (Post Traumatic Stress Disorder), anxiety, and phobias. Its neuronal substrates are key elements to understand the persistent nature of remote memory and open a new perspective to novel therapeutic approaches for human fear-related disorders. While the majority of reports investigate the mechanisms and ensembles of recent fear memory extinction (hours to days following conditioning), only a few refer to the remote time point (i.e. weeks after conditioning), usually describing successful memory extinction. The neuronal correlates of impaired remote fear memory extinction were yet beyond the scope. Here we present selective impairment of contextual remote fear memory extinction in alpha calcium/calmodulin-dependent protein kinase II (αCaMKII) autophosphorylation-limited mice (T286A^+/-^). To map brain regions involved in this phenomenon, we applied screening of c-Fos expression, a neuroplasticity marker, across 23 brain areas following contextual fear conditioning and extinction of recent (1-day old) and remote (30-days old) fear memory in WT and T286A^+/-^ mice. Following impaired remote fear memory extinction in T286A^+/-^ mice, we found upregulated c-Fos expression in the entorhinal cortex (ENT), nucleus reuniens (RE), centromedial (CM), mediodorsal (MD), anterodorsal (AD) thalamic nuclei, and medial septum (MS), compared to WT animals performing normal remote fear memory extinction. Thus our data suggest that αCaMKII-autophosphorylation-dependent c-Fos expression in these areas controls distant contextual fear extinction and may shed light on these brain regions as potential targets for therapeutic strategies against emotional disorders such as PTSD.

## INTRODUCTION

Fear and anxiety are evolutionarily conserved emotions. They aid survival by increasing awareness and enable rapid responses to environmental hazards (LeDoux et al. 2012). Excessive fear and anxiety, on the other hand, are hallmarks of a variety of disabling disorders like phobias or PTSD (Post Traumatic Stress Disorder) that affects millions of people throughout the world (Atwoli et al. 2015; Sareen 2014). A clinical approach to treat PTSD is extinction-based exposure therapy. Extinction learning relies on acquiring new environmental information that suppresses the previously learned fear (Pavlov 1927; Eisenberg et al. 2003; Myers and Davis 2007; Quirk and Mueller 2008; Pape and Pare 2010) as well as some unlearning processes (Khalaf et al. 2018; Bellfy and Kwapis 2020; Dunsmoor et al. 2015; Clem and Schiller 2016). Impaired extinction recall (Milad et al. 2009), and high stability of remote memories (Frankland et al. 2006; Alberini 2011; Gräff et al. 2014; Tsai and Gräff 2014) are proposed as the mechanisms of pathological fear memories conditions observed in PTSD subjects. Still, little is known how distant fear becomes attenuated, thus understanding the neural circuits underlying processing of fear memory, its extinction, and how they drive behavior are especially important in regards to the storage period of the memory.

Hippocampus is a crucial structure for encoding and updating of contextual fear memory (Anagnostaras et al. 1999; Frankland et al. 2007; Varela et al. 2016). The standard model of systems consolidation states that memories are first encoded in the hippocampus and then stored in the cortex (Squire and Bayley 2007). Distribution of the newly gathered information through broadly scattered cortico-hippocampal networks enables its long-term storage and accessibility over time (Frankland et al. 2004; Squire et al. 2015; Vetere et al. 2017; Wheeler et al. 2013).

Long-term memory consolidation involves neuronal remodeling at both the system and synaptic levels (Bailey and Kandel 1993; Dudai 2004; Frankland and Bontempi 2005; Restivo et al. 2009). The basic synaptic processes, that enable storage of behaviorally related information, involve αCaMKII activity (Giese and Mizuno 2013; Lisman et al. 2002; Lisman et al. 2012). αCaMKII is also essential for synaptic plasticity of cortical (Frankland et al. 2001; Hardingham et al. 2003) and subcortical brain areas (Giese et al. 1998; Irvine et al. 2006). Frankland and co-workers reported that αCaMKII^+/-^ heterozygous mice show impaired cortical, but not hippocampal (CA3-CA1), NMDAR-dependent long-term potentiation and their remote long-term contextual fear memory is severely impaired while recent memory is spared (Frankland et al. 2001). Not only αCaMKII activity but also its regulation through autophosphorylation plays an important role in long-term memory formation. αCaMKII autophosphorylation at Threonine 286 (T286) is essential for contextual long-term memory formation and hippocampal LTP (Giese et al. 1998; Irvine et al. 2011; Kimura, Silva, and Ohno 2008). Our group previously showed that the autophosphorylation-deficient mutant mice (αCaMKII^T286A^)(Giese et al. 1998) are impaired in fear memory extinction (Radwanska et al. 2011; Radwanska et al. 2015). However, the studies on the role of autophosphorylation of αCaMKII-T286 in extinction are limited to recent memory extinction paradigms (Kimura et al. 2008; Radwanska et al. 2015), while its role in remote fear memory extinction remains unknown.

In the present study, we tested the role of αCaMKII-T286 in extinction of recent and remote contextual fear and analyzed the brain networks that are activated. To this end, we employed αCaMKII autophosphorylation-deficient heterozygous mice (T286A^+/-^). To reveal the neuronal correlates of remote contextual memory extinction, we employed c-Fos expression mapping. Our results show that T286A^+/-^ mice have exclusively impaired remote contextual fear extinction, while recent memory is spared. Moreover, the impairment is linked with upregulation of fear extinction-induced c-Fos expression in the cortical and thalamic regions in the mutants indicating that these brain areas participate in time- and αCaMKII autophosphorylation-dependent remodeling of contextual fear memory network.

## MATERIALS AND METHODS

### 1. Animals

For this study male mice, weighing 20-30 g were used (n = 3-6 for c-Fos and n = 7 for the behavioral experiment). The animals were maintained on a 12 h light/dark cycle with food and water *ad libitum*. We used the Tg(αCaMKIIT286A)xTg(Thy1-EGFP) mice obtained from crossing αCaMKII autophosphorylation-deficient mutant mice (αCaMKII-T286A) (Giese et al. 1998) and heterozygous of Thy1-GFP M line mice (Thy1-GFP^+/-^) and genotyped as previously described (Feng et al. 2000). Behavioral experiments were conducted in the light phase of the cycle, and mice were 8-10 weeks old at the time of training. All procedures conducted in the study were approved by the 1st Local Ethical Committee, Warsaw, Poland (permission number: 261/2012).

### 2. Contextual fear conditioning

The animals were trained in a conditioning chamber (Med Associates Inc, St Albans, USA) in a soundproof box as previously described (Radwanska et al. 2015). The chamber floor had a stainless steel grid for shock delivery. Prior to the training, the chamber was cleaned with 70% ethanol and a paper towel sprayed with ethanol was placed under the grid floor. In order to camouflage any noise in the behavioral room, background noise was supplied to the chamber by a white noise generator positioned on the side of the soundproof box. On the conditioning day, the mice were brought from the housing room into a holding room where they were allowed to acclimatize for 30 min before training. Next, mice were placed in the chamber and after a 148 s introductory period, a foot shock (2 s, 0.7 mA) was presented. The shock was repeated 5 times, with an inter-trial interval of 90 s. Thirty seconds after the last shock the mouse was returned to its home cage.

Contextual fear memory was tested and extinguished 1 day (recent, also called Rec) or 30 days (remote also called Rem) after training by re-exposing the mouse to the conditioning chamber for 20 minutes (extinction session), and they were sacrificed 90 min after the beginning of the extinction session. Control mice underwent the same procedure but without shock presentation (CTX) or were anesthetized twenty-six hours after training (without extinction session) (5US), or were taken directly from their home cages (Naive) (Figure 1C). Freezing behavior and locomotor activity were scored by the experimenter blind to the mice genotype from the movies recorded by a video camera placed on the door of the sound attenuating box and measured automatically by Video Freeze® software (Med Associates Inc, St Albans, USA).

**Figure 1.**
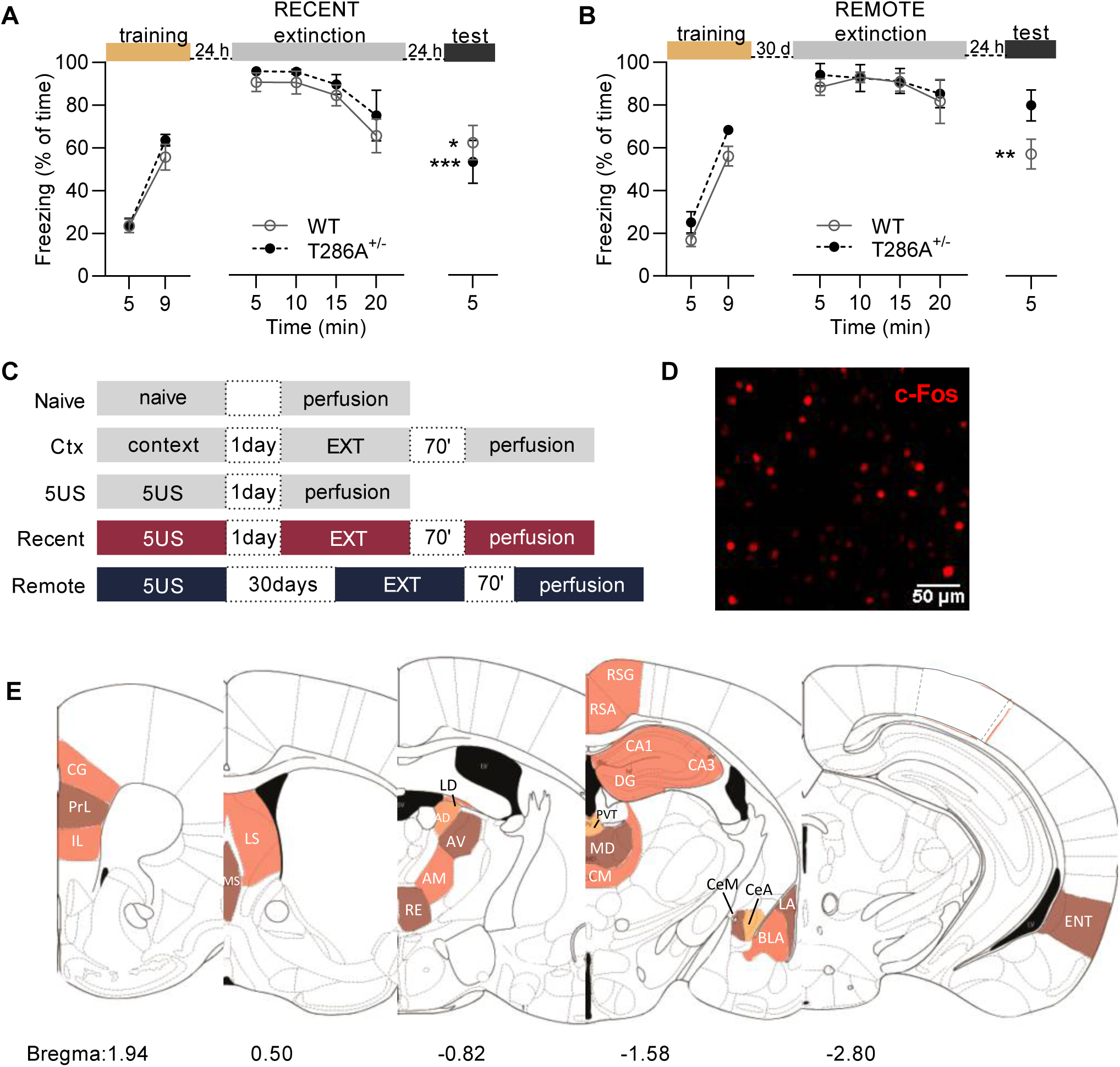
Consolidation of remote fear extinction memory is impaired in T286A^+/-^ mice. **(A-B)** Experimental timeline and freezing levels of WT (open circle), and T286A^+/-^ (black, full circle) mice during fear conditioning, recent **(A)** or remote **(B)** fear extinction session and retention test of fear extinction memory. The WT mice, as well as T286A^+/-^ exhibited normal contextual recent fear extinction, as they freeze significantly less during the test, compared to the first 5 minutes of extinction session. However, T286A^+/-^ mice showed impaired remote fear extinction consolidation compared to WT littermates. Two-way ANOVA, and Tukey’s *post-hoc* test were used; data with SEM * p < 0.05, **p < 0.01 **, *p < 0.001. **(C)** An experimental design for c-Fos expression analysis. **(D)** An example of a microphotograph of immunolabeled c-Fos protein. **(E)** An overview of analyzed brain structures.

### 3. c-Fos immunoreactivity

The mice were deeply anesthetized with sodium pentobarbital (50 mg/kg intraperitoneal (i.p.)) and perfused transcardially with 20 ml of PBS (Phosphate-buffered saline; POCH, Poland), followed by 50 ml of 4% paraformaldehyde (PFA; Sigma-Aldrich, Poland) in 0.1 M PBS, pH 7.5. Brains were stored in the same solution for 24 h at 4℃. Next, the brains were transferred to the solution of 30% sucrose in 0.1 M PBS (pH 7.5) and stored at 4℃ for 72h. Brains were frozen at -20℃ and coronal sections (40 µm) cut in the frontal plane on a cryostat microtome (YD-1900, Leica). Coronal slices were stored in -20℃ in PBSAF (PBS, 15% sucrose, 30% ethylene glycol, 0.05% NaN3). Every sixth section through the whole brain was used for immunostaining. Sections were washed three times for 6 min in PBS, followed by incubation in blocking solution (5% NDS; Jackson Immuno Research/0,3% Triton X-100; Sigma Aldrich) in PBS for 2h at room temperature (RT). Next, sections were incubated with c-Fos polyclonal antibody (sc-52; Santa Cruz Biotechnology, 1:1000) for 12h in 37℃, washed three times in TBS (0,3% Triton X-100 in PBS) and incubated with secondary antibody (1:500) conjugated with Alexa Fluor 594 (Invitrogen A-11062) for 2h at room temperature. After incubation with the secondary antibody slices were washed three times for 6 min in PBS and mounted on microscopic slides covered with mounting dye with DAPI (Invitrogen, 00-4959-52).

### 4. Immediate Early Gene (IEG) quantification

c-Fos expression was analyzed in 23 brain regions, where c-Fos positive cells were counted with 10x magnification, in the red spectrum (568 nm laser excitation) under a confocal microscope (SP5, Leica). Each photomicrograph was followed with image acquisition in a blue-green spectrum (488 nm laser excitation) that highlighted major structures in the brain of mice with Thy1-GFP background. Cells were considered positive for c-Fos immunoreactivity if the nucleus was the appropriate size (area ranging from 5 to 160 µm^2^) and was distinct from the background. For each photomicrograph converted into an 8-bit grayscale, the same threshold was used and selected areas of interest were measured using ImageJ software (National Institute of Health, Bethesda, MD). The densities of the nuclei were counted (by an experimenter blind to the experimental condition) as a number of immunostained nuclei within the analyzed region, divided by the area of that region (measured on the picture and expressed in pixels). The position of the analyzed brain regions was determined according to the atlas of the mouse brain (Franklin and Paxinos 2019).

### 5. Statistics

For statistical analysis one-way or two-way analysis of variance (ANOVA) with Sidak and Tukey’s *post-hoc* test for multiple comparisons was used when appropriate. Differences between the experimental groups were considered significant if *p*□<□0.05. Statistical tests and the results are detailed in the text, Sup. Figure 1. and figure legends. All data were analyzed with GraphPad Prism 8 software for Windows (GraphPad Software, San Diego, California USA, www.graphpad.com). Anatomical cases were excluded if they had significant damage to the brain tissue.

## RESULTS

### 1. αCaMKII T286A^+/-^ mutation results in contextual remote fear memory extinction deficits

Contextual fear conditioning training was used to test the formation and extinction of recent (1 day) and remote (30 days) memory in mice. Mice (n = 7 per experimental group) were trained in a novel context according to a previously published protocol (5 × 0.7 mA; 2 s) (Radwanska et al. 2015). After 1 or 30 days, animals underwent a fear extinction session (EXT - 20 minutes of exposure to the experimental context without US presentation) to extinguish recent or remote contextual fear memory. On the following day (day 3 or 32), the fear extinction memory was tested in the same context for 5 minutes (TEST) **(Figure 1A**).

Contextual fear conditioning induced the formation of robust recent and remote contextual fear memory both in wild-type (WT) mice and heterozygotes of IZCaMKII autophosphorylation-deficient mutant mice (T286A^+/-^), measured as freezing in the conditioned context (day 2 and day 31, respectively) (**Figure 1A-B**). Both WT mice and T286A^+/-^ heterozygous mutants presented a significant decrease in levels of contextual freezing during the recent fear extinction test (day 3), as compared to the beginning of extinction session (day 2) (Two-way ANOVA showed only a significant effect of time point [F_(6, 63)_ = 31.6, p < 0.001] on freezing levels), confirmed by *post hoc* Tukey’s test (WT: p = 0.05 and T286A^+/-^: p < 0.001). A similar decrease during remote fear extinction testing (day 31 vs. 32) was observed (Two-way ANOVA showed a significant effect of time point [F_(6, 77)_ = 40.76, p < 0.001] and training [F_(1, 77)_ = 6.219, p = 0.01] on freezing levels) only in the WT group (p = 0.008), but not in T286A^+/-^ mice (p = 0.82), where no significant change in the level of freezing was observed between the sessions (**Figure 1B**). This observation shows impaired remote contextual fear extinction memories in T286A^+/-^ mice.

### 2. The effect of T286A^+/-^ mutation on fear extinction-induced c-Fos expression in the dorsal hippocampus

To examine the activity of brain structures involved in impaired remote fear memory extinction of T286A^+/-^ mice, we analyzed c-Fos expression as a proxy of neuronal activity (Sagar et al. 1988; Guzowski et al. 2005). c-Fos was analyzed in five experimental groups (animals per group: n = 3-6) (**Figure 1C**). c-Fos-positive cellular nuclei have been counted in the dorsal hippocampus, amygdala, cortex, thalamus, and septum, a total of 23 brain regions (**Figure 1F, Supplementary. Figure 1, Supplementary Table 1**); the two-way ANOVA F-statistic and *post-hoc* tests collected in the table for each structure can be found in (**Supplementary Table 2**).

Contextual fear conditioning and extinction have been shown to be hippocampus-dependent (Bissiere et al. 2011; LeDoux and Phillips 1992; Maren and Holt 2000; Maren and Quirk 2004). In the CA1 field of hippocampus we observed significant effects of genotype on c-Fos expression, however, the post hoc tests revealed no significant differences in c-Fos counts between the experimental groups (**Figure 2A**). The CA3 showed significant effects of genotype and training for c-Fos expression, but similarly to CA1, there were no significant differences in the c-Fos signal (**Figure 2B**). Only the WT Naive group has more c-Fos positive cells than T286A^+/-^ (p = 0.024). In the DG we have detected a significant effect of genotype and training on c-Fos expression. WT mice demonstrated increased c-Fos expression in Context group (p = 0.01) compared to the Naive group, but when compared to 5US group similar increase was noticed in Context group (p < 0.001), Rec (p < 0.001) and Rem (p < 0.03) extinction (**Figure 2C**). In the hippocampus of T286A^+/-^ mice we found no significant changes in the c-Fos signal between experimental groups.

**Figure 2.**
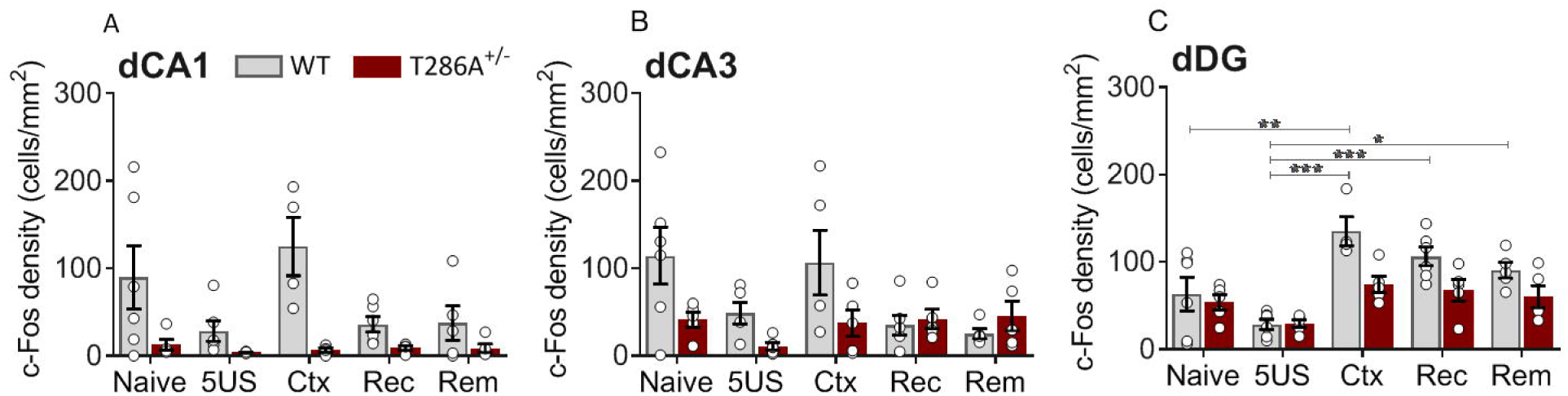
A c-Fos expression in the dorsal hippocampus fields CA1, CA3, and DG do not differ between WT and T286A^+/-^ mice subjected to contextual fear conditioning and extinction. Expression of c-Fos in the dorsal hippocampus. Gray bars represent WT, control animals, while red bars represent T286A^+/-^ αCaMKII mice. **(A)** In the dCA1, no significant differences were observed in c-Fos expression, in both WT (n=5-6) and T286A^+/-^ (n=4-6) mice. **(B)** Naive group of T286A^+/-^ (n=5-6) animals exhibit diminished c-Fos levels in the dCA3 field of the hippocampus compared to WT (n=5-6) controls. **(C)** In the dDG, there were no significant differences in the c-Fos levels between WT (n=4-6) and T286A^+/-^ (n=5-6) mice across all training conditions. Means ± SEM are shown; Two-way ANOVA with Sidak *post hoc* test, gray * indicates significant difference between conditions within the WT genotype, while red T286A^+/-^; # indicates a significant difference between genotypes, within the condition. * p < 0.05, ** p < 0.01, *** p < 0.001.

### 3. The effect of T286A^+/-^ mutation on fear extinction-induced c-Fos expression in the amygdala

Amygdala is involved in the encoding of contextual fear (Herry et al. 2010; LeDoux and Phillips 1992; Lee et al. 2013; Orsini and Maren 2012; Ehrlich and Josselyn 2016) and is also known to be involved in recent (Ehrlich et al. 2009; Herry and Mons 2004; Hobin et al. 2003) and remote fear extinction (Cambiaghi et al. 2016; Gale et al. 2004; Silva et al. 2018; da Silva et al. 2020). Therefore, we measured c-Fos densities in basolateral (BLA), lateral (LA), central (CeA - consisting of capsular and lateral division that were analyzed together), and centromedial (CeM) nuclei of amygdala (**Figure 3**).

**Figure 3.**
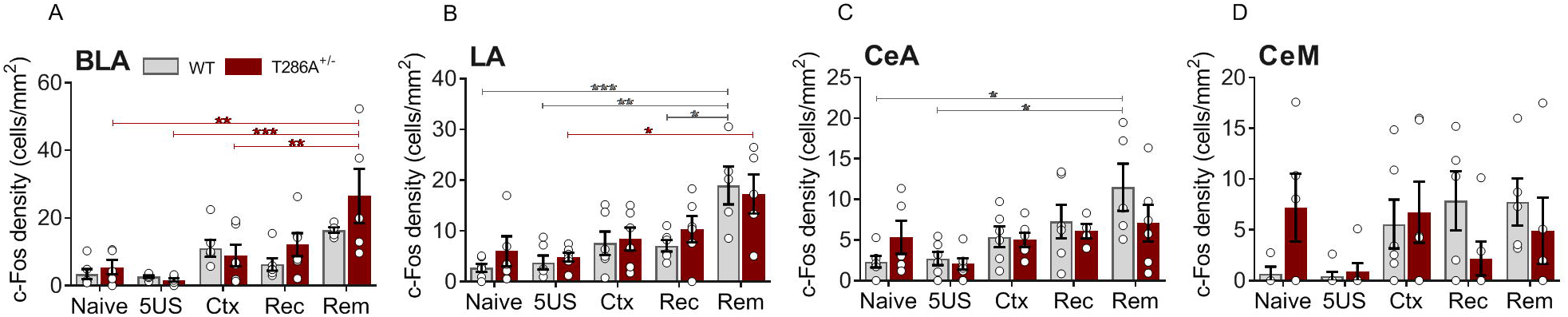
Regional activation of amygdalar structures did not differ between both genotypes, WT and T286A^+/-^ following extinction learning. c-Fos density in amygdala. Gray bars represent WT, control animals, while red bars represent T286A^+/-^ IZCaMKII mice. **(A-D)** No changes of c-Fos expression were observed between T286A^+/-^ and WT animals in any of the subregions of amygdala. Means ± SEM are shown. For WT (n=5-6 in BLA, LA and CeA, n=4-6 in CeM). For T286A^+/-^(n=5-6 in BLA and LA, n=4-6 in CeA, n=5-6 in CeM). Two-way ANOVA with Sidak *post hoc* test, gray * indicates significant difference between conditions within the WT genotype, while red T286A^+/-^; * or p < 0.05, ** p < 0.01, *** p < 0.001.

BLA showed a significant effect of training, but not genotype, on c-Fos expression. In T286A^+/-^ mice c-Fos density was elevated in BLA following remote extinction when compared to Naive (p = 0.001), 5US (p < 0.001) and Ctx group (p < 0.01) (**Figure 3A**). In the LA, a significant effect of training on c-Fos was observed. Expression of c-Fos, when compared to the 5US group, was elevated after remote extinction in both genotypes WT (p = 0.002) and T286A^+/-^ (p = 0.021) mice. Whereas only WT animals presented increased c-Fos density in the Rem group compared to Naive (p < 0.001) and Rec (p = 0.034) of the same genotype (**Figure 3B**). In the CeA we have noticed a significant training effect on c-Fos expression. WT animals expressed more c-Fos following remote extinction than Naive animals (p < 0.011) and 5US (p = 0.019) group of the same genotype (**Figure 3C**). Such differences were not observed in the mutants. The analysis of c-Fos expression in CeM showed neither significant genotype × training interaction nor effects of genotype or condition (**Figure 3D**). Thus T286A^+/-^ and WT animals exhibited similar c-Fos expression patterns in BLA, LA, CeA and CeM across all training conditions. This confirms that the amygdala is an important component of the fear extinction memory circuit, however, its activity during the training is not regulated by αCaMKII.

### 4. The effect of T286A^+/-^ mutation on fear extinction-induced c-Fos expression in the cortex

The medial prefrontal cortex (mPFC), including both prelimbic (PrL) and infralimbic (IL) areas, is implicated in fear conditioning and extinction (Awad et al. 2015; Do-Monte et al. 2015; Frankland et al. 2004; Giustino and Maren 2015; Peters et al. 2009; Ramanathan et al. 2018; Wheeler et al. 2013). The cingulate cortex (CG) is activated by remote contextual fear memory, as shown by c-Fos and Zif268 imaging (Frankland et al. 2004), moreover, the persistence of reactivated fear memory becomes CG-dependent with time (da Silva et al. 2020). Two subdivisions of retrosplenial cortex, namely granular (RSG) and agranular (RSA), as well as entorhinal cortex (ENT) were also selected since they communicate directly with the hippocampus (Baldi and Bucherelli 2014; 2015; Corcoran et al. 2011; Suh et al. 2011).

In the PrL there was a significant genotype × training interaction and significant effect of training, but not genotype, on c-Fos expression. WT mice had similar levels of c-Fos-positive nuclei density in every condition, while significantly higher levels were observed in the remote extinction of T286A^+/-^ mice compared to the Rec (p = 0.01), Ctx (p = 0.007), 5US (p < 0.001) and Naive (p = 0.003) groups of the same genotype. The *post hoc* tests found no significant differences between T286A heterozygotes and WT in c-Fos levels, in all five conditions (**Figure 4A**). In the IL, a significant effect of training on c-Fos levels was observed. Compared to Naive group of WT animals elevated expression of c-Fos was found in Ctx (p = 0.030), Rec (p = 0.036) and Rem (p = 0.010) groups, while comparing to 5US only in Ctx (p = 0.041) and Rem (p = 0.014) of the same genotype. In T286A^+/-^ mice, increased levels of c-Fos expression were noted in the animals following context exposure (p = 0.024 compared to 5US group) and after remote extinction (p = 0.024 compared to Naive and p < 0.001 compared to 5US). Both WT and T286A^+/-^ animals expressed similar levels of c-Fos across all five training conditions (**Figure 4B**). In the CG we also found a significant effect of training on c-Fos density. In WT animals an increased c-Fos expression was found in the Ctx group when compared to Naive (p = 0.035) and 5US (p = 0.020). T286A^+/-^ mice in turn showed more c-Fos positive cells in the Rec (p = 0.031) and Rem extinction groups (p = 0.005) when compared to the 5US group (**Figure 4C**). In the RSA there was neither significant genotype × training interaction nor effect of genotype or training on c-Fos expression (**Figure 4D**). However, in the RSG, a significant effect of training on c-Fos density was found; following remote extinction (p = 0.036) the levels of c-Fos expression were higher compared to the Naive group. No differences between WT and T286A^+/-^ mice were detected in any of the experimental groups (**Figure 4E**). There was a significant genotype × training interaction, effect of training and genotype on c-Fos levels in the ENT. T286A^+/-^, but not WT animals, exhibited significant increase c-Fos densities following remote extinction when compared to Naive (p < 0.001), 5US (p < 0.001), Ctx (p = 0.001) and Rec (p = 0.001) groups. We also observed more c-Fos-positive cells in T286A^+/-^ mice after remote extinction, compared to WT littermates (p = 0.011) (**Figure 4F**).

**Figure 4.**
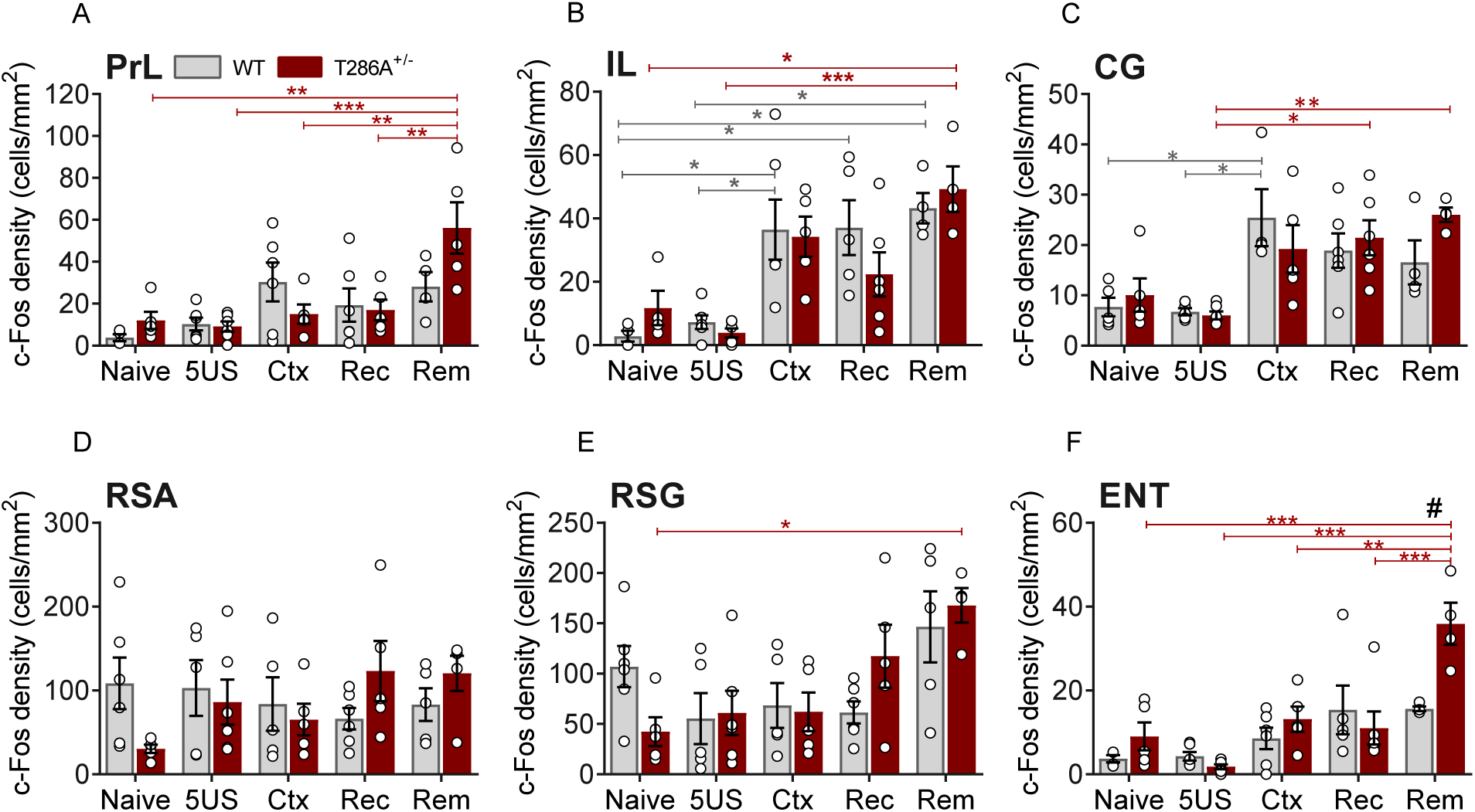
T286A^+/-^ αCaMKII but not WT mice display hyperactivation in the ENT cortex following remote fear memory extinction. Expression of c-Fos in cortical structures. Gray bars represent WT, control animals, while red bars represent T286A^+/-^ αCaMKII mice. **(A-E)** In the PrL (n=4-6 for both WT and T286A^+/-^), IL (n=4-6 for both WT and T286A^+/-^), CG (n=4-6 for both WT and T286A^+/-^), RSA (n=5-6 for both WT and T286A^+/-^) and RSG (n=5-6 for WT and n=4-6 for T286A^+/-^) the c-Fos expression remained similar between WT and T286A^+/-^ animals in all five conditions. **(F)** In the ENT, T286A^+/-^ (n=5-6) mice exhibit elevated c-Fos density following remote extinction in contrast to their WT (n=4-6) littermates. Means ± SEM are shown. Two-way ANOVA with Sidak *post hoc* test, gray * indicates significant difference between conditions within the WT genotype, while red in T286A^+/-^; # indicates a significant difference between genotypes, within the condition. * or # p < 0.05, ** p < 0.01, *** p < 0.001.

### 5. The effect of T286A^+/-^ mutation on fear extinction-induced c-Fos expression in the thalamus

Our data indicate that the PrL and ENT are hyperactivated by remote fear attenuation in the T286A^+/-^ mice, as compared to other training groups. As these brain structures are directly interconnected with hippocampus and amygdala (Do-Monte et al. 2015; Ehrlich et al. 2009; Hefner et al. 2008; Herry et al. 2010; Milad and Quirk 2012; Vertes et al. 2015) or by surpassing the thalamic and septal nuclei (Aggleton et al. 2016; Do-Monte et al. 2015; Krettek and Price 1977; Unal et al. 2015) we chose a group of anterior (anterodorsal - AD, anteromedial - AM, anteroventral - AV, laterodorsal - LD) and midline (centromedial - CM, mediodorsal - MD, paraventricular - PVT, nucleus reuniens - RE) thalamic nuclei together with medial (MS) and lateral (LS) septal nuclei to examine their c-Fos expression **(Figure 5)** Moreover, contemporary studies implicate PVT (Do-Monte et al. 2015; Penzo et al. 2015), RE (Ramanathan et al. 2018; Silva et al. 2018; Vetere et al. 2017; Wheeler et al. 2013; Silva et al. 2021), MD (Bradfield et al. 2013; Lee et al. 2012; Li et al. 2004) and LS (Vetere et al. 2017) in threat memory processing but their role in remote fear memory attenuation, remains elusive.

**Figure 5.**
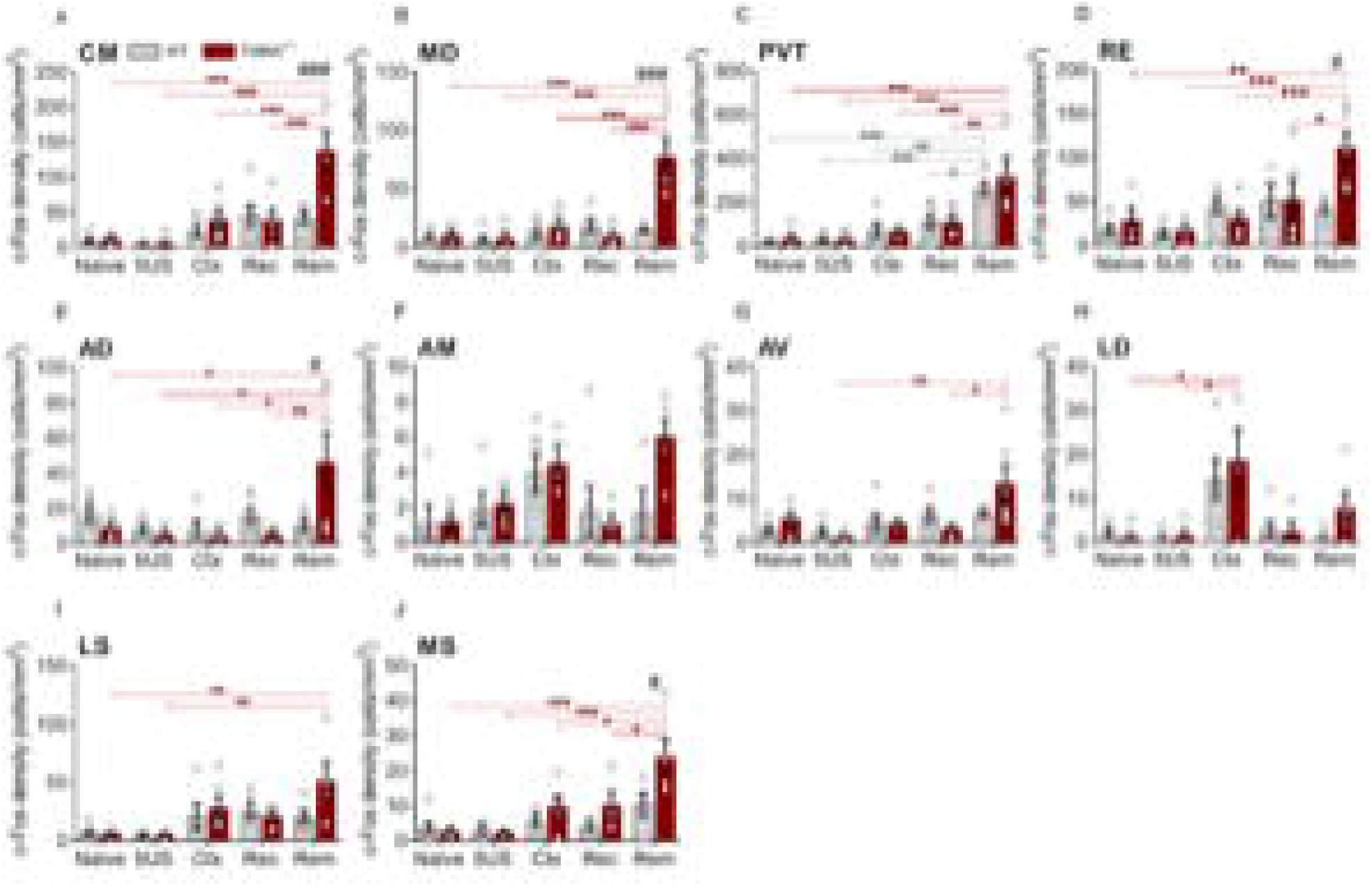
Vast activation of CM, MD, RE, AD, and MS nuclei following remote extinction in T286A^+/-^ αCaMKII mice, but not WT. Expression of c-Fos in midline thalamus, anterior thalamus, and septum. Gray bars represent WT, control animals, while red bars represent T286A^+/-^ αCaMKII mice. **(A-B)** Increased c-Fos density was observed in T286A^+/-^ compared to WT, following remote extinction in CM and MD (n=5-6 for WT and n=4-6 for T286A^+/-^ mice in both regions). **(C)** In the PVT, no differences between WT (n=5-6) and T286A^+/-^ (n=4-6) mice were noted in any of the five conditions. **(D-E)** T286A^+/-^ animals that underwent remote extinction procedure exhibited significantly elevated levels of c-Fos in RE (n=3-6 for WT and n=4-6 for T286A^+/-^) and AD (n=5-6 for WT and n=3-6 for T286A^+/-^) as compared to their WT littermates. **(F-I)** In the AM (n=4-6 for WT and n=3-6 for T286A^+/-^), AV (n=4-6 for WT and n=3-6 for T286A^+/-^), LD (n=4-6 for WT and n=4-5 for T286A^+/-^) and LS (n=5-6 for WT and n=4-6 for T286A^+/-^), the WT and T286A^+/-^ mice have comparable c-Fos densities across all five conditions. **(J)** Again, in the MS, the T286A^+/-^ (n=3-6) mice have significantly more c-Fos-positive cells than WT (n=3-6) animals following remote extinction. Means ± SEM are shown. Two-way ANOVA with Sidak *post hoc* test, gray * indicates significant difference between conditions within the WT genotype, while red T286A^+/-^; # indicates a significant difference between genotypes, within the condition. * or # p < 0.05, ** p < 0.01, *** or ### p < 0.001.

In the CM we observed a significant interaction between training and genotype, as well as a significant effect of genotype and condition on c-Fos levels. The *post hoc* Sidak tests found that T286A^+/-^ mice of Rem group showed elevated c-Fos expression compared to that in Naive, 5US, Ctx, and Rec (p < 0.001 equally). T286A^+/-^ mice have a significantly higher expression of c-Fos following remote extinction than the control WT animals (p < 0.001) **(Figure 5A)**. Similar to the CM, there was a genotype × training interaction, the effect of genotype and training on c-Fos expression in MD. The remote extinction group of T286A^+/-^ mice had increased density of c-Fos compared to Naive, 5US, Ctx, and Rec (p < 0.001; *p-value* equal to every comparison) of the same genotype. Moreover, this group also had higher c-Fos levels than WT littermates sacrificed after a remote extinction (p < 0.001) **(Figure 5B)**. In the PVT there was a significant effect of training on c-Fos expression. Both WT and T286A^+/-^ mice demonstrated increased c-Fos counts after remote extinction: in WT compared to Naive (p < 0.001), 5US (p < 0.001), Ctx (p = 0.002) and Rec (p = 0.042) of the same genotype; in T286A^+/-^ analogously Naive (p < 0.001), 5US (p < 0.001), Ctx (p < 0.001), and Rec (p = 0.004). The *post hoc* Sidak test revealed no significant difference in c-Fos density between Rem extinction groups of WT and T286A^+/-^ mice **(Figure 5C)**. In the RE we observed a significant genotype × training interaction and the effect of training on c-Fos levels. *Post hoc* test showed, however, that only the T286A^+/-^ animals following remote extinction had raised c-Fos counts compared to the Naive (p = 0.002), 5US (p < 0.001), Ctx (p < 0.001) and Rec (p = 0.041) groups of the same genotype. Moreover, following the remote extinction training T286A^+/-^ mice showed significantly more c-Fos positive cells than their WT littermates (p = 0.016) **(Figure 5D)**. Within the anterior division of the thalamus, there was a significant interaction between the condition and training and the effect of training on the expression of c-Fos in AD. Again, significantly more c-Fos-positive cells was detected after remote extinction, compared to Naive (p = 0.033), 5US (p = 0.020), Ctx (p = 0.049) and Rec (p = 0.010) groups, exclusively in T286A^+/-^ mice. Additionally, the Rem group of T286A^+/-^ mice presented higher levels of c-Fos than Rem WT animals (p = 0.049) **(Figure 5E)**. In the AM we found a significant effect of training on c-Fos density. Nevertheless, significant differences were found neither within nor between the genotypes in all five training groups **(Figure 5F)**. In the AV there was a significant effect of training on c-Fos levels. *Post hoc* Sidak tests revealed elevated c-Fos levels in the Rem group of T286A^+/-^ mice in contrast to the 5US (p = 0.005) and Rec (p = 0.018) conditions, however, there were no significant differences in c-Fos density between the genotypes **(Figure 5G)**. In the LD we have observed a significant effect of training on c-Fos counts, but no effect of the genotype. The T286A^+/-^ animals showed increased c-Fos counts following context exposure as compared to the control groups: Naive (p = 0.017) and 5US (p = 0.038) **(Figure 5H)**. There was a significant effect of training, but not genotype, on c-Fos expression in LS. However, the *post hoc* tests revealed that only T286A^+/-^mice expressed significantly more c-Fos after remote extinction than the Naive (p = 0.009) or 5US (p = 0.006) mice of the same genotype **(Figure 5I)**. In the MS, there were statistically significant effects of training and genotype on c-Fos levels. Only T286A^+/-^ mice had higher c-Fos density in the remote extinction group as compared to the Naive (p < 0.001), 5US (p < 0.001), Ctx (p = 0.016) and Rec (p = 0.048) control groups. There were also significantly more c-Fos positive cells in T286A^+/-^ MS than in the WT following remote extinction (p = 0.030) **(Figure 5J)**.

## DISCUSSION

In the current study, we analyzed neuronal correlates of remote contextual fear memory extinction and the role of autophosphorylation of IZCaMKII in this process. We have shown for the first time that the T286A^+/-^ mice exhibit impaired extinction of remote contextual fear memory, with spared extinction of recent contextual fear. Our results demonstrate that remote fear memory extinction failure was associated with upregulated c-Fos levels in the ENT, RE, CM, MD, AD, and MS. c-Fos immunomapping have been previously used to analyze neuronal substrates of many aspects of contextual fear memory processing, including fear memory encoding (Park and Chung 2019; Lin et al. 2018; Frankland et al. 2004), recall (Wheeler et al. 2013; Hall et al. 2001; Conejo et al. 2007), recent extinction (Park and Chung 2019; Knox et al. 2016), remote extinction (Silva et al. 2018) and even extinction deficits (Park and Chung 2019; Hefner et al. 2008; Talukdar et al. 2018). Nevertheless, none of these studies investigated the role of autophosphorylation of IZCaMKII in remote fear memory extinction. None of them, also have mapped the activity of brain regions associated with remote fear extinction-deficits that resemble clinically observed emotional disorders, such as PTSD (Milad et al. 2009).

T286A mutants have severely impaired contextual fear memory after a single conditioning trial, but after a massed training protocol, T286A mutants can form contextual recent (1 d) and remote (30 d) long-term memory (Irvine et al. 2005; Irvine et al. 2011). This fact overlaps with our findings on T286A^+/-^ mutants - after intensive training they expressed both, recent and remote fear memory at the levels of WT mice. Moreover, it has been shown that extinction of recent contextual fear is impaired in T286A (Radwanska et al. 2011; Radwanska et al. 2015) and in T286A^+/-^ mutant mice (Kimura et al. 2008). In the present study, we show for the first time that T286A^+/-^ mice exhibit intact recent but impaired remote fear memory extinction. In our experiments, WT and T286A^+/-^ mice learned similarly during recent extinction session, while Kimura *et al*. (2008) observed better extinction performance in WT mice, in an adequate paradigm (24 h after conditioning). This discrepancy likely results from the differences in the strength of fear conditioning training (1 vs. 5US). Thus our new data and the review of the literature indicate that autophosphorylation of IZCaMKII affects persistence and stability of contextual fear memory and that deficits in IZCaMKII activity can be overcome by the intensity of the training (Irvine et al. 2005; Irvine et al. 2011).

In order to characterize brain circuits activated by impaired extinction, we used c-Fos immunostaining to map brain regions related to contextual fear, and extinction learning.

The hippocampus is involved in context-dependent learning and fear memory (Kim and Fanselow 1992). Even in the absence of □CaMKII autophosphorylation, contextual fear conditioning is hippocampus-dependent (Irvine et al. 2011). Studies on αCaMKII^+/-^ mutants indicate that c-Fos expression in the CA1 and/or CA3 areas of the hippocampus remains unchanged after fear conditioning, recent or recall of contextual fear, as compared to WT mice (Frankland et al. 2004; Yamasaki et al. 2008). The remote recall of contextual fear downregulates c-Fos in the hippocampus of WT (Gräff et al. 2014; Wheeler et al. 2013), but not in αCaMKII^+/-^ mice (Frankland et al. 2004). Contrary to our observation, recent extinction of contextual fear enhances c-Fos expression in the hippocampus of WT as compared to the Naive group of mice (Park and Chung 2019). This is possibly due to the difference in the basal levels of c-Fos in Naive groups or, again fear conditioning protocol (3 vs. 5 US). Noteworthy, we also observed an upregulation of c-Fos signals in DG, in CTX, Rec, and Rem group of WT mice compared to 5US condition, which confirms the involvement of DG in context perception and extinction learning (Khalaf et al. 2018; McHugh et al. 2007; Besnard and Sahay 2016). The remote extinction, although does not change c-Fos expression in dorsal CA1, CA3, and DG (Silva et al. Gräff 2018), which is in agreement with our results, except for DG. However, to our knowledge, there is no study examining IEG expression in recent or remote fear extinction memories of the T286A^+/-^ mice. Our analysis of c-Fos activation in the hippocampus revealed a significant effect of genotype on c-Fos levels expression in the DG, CA1, and CA3 areas of the hippocampus, however no differences between WT and T286A^+/-^ mice across all tested conditions were significant.

Nevertheless, it has been also proposed that the hippocampus remodel remote memories in the absence of the original trace, an idea coined as a “scene construction theory” (for review see: Barry and Maguire 2019), thus unchanged neural activation in the hippocampus during remote fear extinction may not exclude its potential involvement in remote memory processing (see: Goshen et al. 2011; Tayler et al. 2013). Complementary to this, a new study revealed that the CA3 field of hippocampus plays a modulatory role of remote, but not recent recall of fear memory *via* long-range inhibitory pathway to anterodorsal thalamus nucleus (Vetere et al. 2021). Thus, it appeared plausible that similar mechanisms may be implicated in other hippocampus-originating inhibitory long-range connections, as some examples of such circuits have also been previously reported in cortical (Melzer et al. 2012; Jinno et al. 2007; Yamawaki et al. 2019), septal (Takács et al. 2015; Tóth and Freund 1992), and amygdalar nuclei (Lübkemann et al. 2015). However, such discrete, usually sparse projection could be missed with the c-Fos labeling approach that we used here.

Another dominant brain hub, functionally interconnected with hippocampus and implicated in fear learning and its extinction is amygdala (Herry et al. 2010; LeDoux and Phillips 1992; Lee et al. 2013; Orsini and Maren 2012; Ehrlich et al. 2009; Ehrlich and Josselyn 2016; Herry and Mons 2004; Hobin et al. 2003; Cambiaghi et al. 2016; Gale et al. 2004; Silva et al. 2018).

Successful extinction of recent contextual fear does not specifically elevate c-Fos expression in amygdala, which is in line with our results (Park and Chung 2019). Our c-Fos analysis shows a significant effect of training on c-Fos expression in LA, CeA, and BLA, and enhanced c-Fos expression in LA, CeA, and CeM of WT mice following a remote fear extinction session, supporting their role in this phenomenon (**Figure 3**). In contrast, the BLA exhibited no differences in c-Fos levels between conditions in WT mice, but a significantly stronger activation during remote extinction was detected in T286A^+/-^ mice (**Figure 3A**). This stands in opposition to the results obtained by Silva et al. (2018), where WT mice reflected higher c-Fos levels upon successful remote extinction of contextual fear, although authors analyzed BLA complex (basolateral and lateral nuclei together), while we have separated those structures. Nevertheless, our results suggest that αCaMKII activity in BLA is involved in, but not critical for remote fear memory processing. There are, however, no unidirectional conclusions on whether the αCaMKII activity is essential for amygdala-dependent learning, moreover, the role of αCaMKII in amygdala has not been studied extensively. For example, one study shows that inhibition of αCaMKII activity in LA by KN-62 leads to impairment of fear memory acquisition (Rodrigues et al. 2004). However, the c-Fos expression in BLA of αCaMKII^+/-^ mice was similar to WT littermates following contextual fear learning (Yamasaki et al. 2008). Our pairwise comparisons of c-Fos expression showed no significant differences in c-Fos activation between T286A^+/-^ and WT mice across all conditions, in any of the regions of amygdala (**Figure 3**). Therefore, our observations provide another piece of evidence in understanding the role of the αCaMKII in amygdala-dependent learning and suggest that αCaMKII activity in the amygdala is unlikely to be responsible for fear extinction failure.

The medial prefrontal cortex projections originating from the hippocampus are implicated in fear memory extinction (Knapska et al. 2012; Giustino and Maren 2015). Several reports found the retrosplenial and entorhinal cortices to be involved in both fear extinction and contextual learning in remote time-points (Corcoran et al. 2013; Silva et al. 2018; Baldi and Bucherelli 2014). In the current study, we have noticed a lack of significant changes in c-Fos signal between T286A^+/-^ and WT groups in any of the conditions in PrL, IL, CG divisions of mPFC, and retrosplenial cortex (**Figure 4**). It is likely that IL and CG are not directly implicated in context-dependent extinction learning, as we observed no differences in c-Fos expression between extinction and context groups, regardless of high c-Fos activation, confirming the proposed role of these structures in recognizing the contextual, rather than an emotional component of fear memory (Silva et al. 2018).

Interestingly, the PrL, RSG, and ENT exhibited c-Fos hyperexpression following impaired extinction. Nevertheless, only ENT activity was significantly higher in T286A^+/-^ than in WT mice (**Figure 4**). High upregulation of PrL activity follows an analogical activation pattern found in BLA, which is unique for impaired extinction in T286A^+/-^ mice. This confirms the previous observation that rats with poor extinction memory showed higher unit activity of PrL than animals with good extinction memory (Burgos-Robles et al. 2009). PrL and BLA are anatomically and functionally interconnected (Vertes 2004; Hoover and Vertes 2007). The PrL is responsible for eliciting fear (Milad and Quirk 2012; Peters et al. 2009; Knapska and Maren 2009; Quirk and Beer 2006; Sierra-Mercado et al. 2011), while BLA is implicated in extinction memory encoding and consolidation (Herry and Mons 2004; Zhang et al. 2020), but also in regulating the expression of IEGs in the hippocampus during contextual fear memory acquisition (Huff et al. 2006; Tronson et al. 2012). Additionally, the excitatory PrL-BLA pathway is activated when animals encounter the fearful context at a remote time point to promote a high fear state (Arruda-Carvalho and Clem 2014), and is presumably suppressed when fear is attenuated (Cho et al. 2013; Laricchiuta et al. 2021). Consequently, the PrL-BLA connection is enhanced when the extinction of fear is impaired (Park and Chung 2020). Poor extinction retention also correlates with upregulated c-Fos in these structures (Stafford et al. 2013). These findings are further supported by our observation, suggesting that PrL is likely working in concert with BLA upon extinction failure of remote contextual fear, although this mechanism is probably not αCaMKII autophosphorylation-dependent.

Finally, we show that the entorhinal cortex is actively engaged when a mouse is unable to extinguish remote contextual fear. Three lines of evidence support this hypothesis. Firstly, inhibition of the BLA-ENT impairs the acquisition of contextual fear memories (Sparta et al. 2014). Secondly, blocking the ENT activity deteriorates extinction of contextual fear (Baldi and Bucherelli 2014). And finally, blocking CaMKII activity in this structure prevents extinction of inhibitory avoidance memory (Bevilaqua et al. 2006), suggesting that the ENT is an element of the neuronal circuit of defective extinction and plausibly acts as a discriminator of the emotional component of contextual fear.

A considerably large extent of evidence supporting thalamic nuclei function in fear learning and its extinction has been provided for PVT (Do-Monte et al. 2015; Chen and Bi 2019; Penzo et al. 2015; Barson, Mack, and Gao 2020, RE (Ramanathan and Maren 2019; Silva et al. 2018; Wheeler et al. 2013; Vetere et al. 2017; Troyner and Bertoglio 2021, Silva et al. 2021), MD (Corcoran et al. 2016; Herry and Garcia 2002; Li et al. 2004; Lee and Shin 2016) and CM (Furlong, Richardson, and Mcnally 2016). Relatively less is known about anterior thalamic nuclei, however their connections with hippocampus, anterior cingulate and retrosplenial cortices suggest that they might be involved in remote and contextual memory processing (Jankowski et al. 2013), but also in fear and extinction (Corcoran et al. 2016; Marchand et al. 2014; Dupire et al. 2013; Jenkins et al. 2002; Vetere et al. 2021). Anteriorly to the thalamus is septum, whose principal role is to control theta rhythms of the hippocampus (for review: Buzsáki 2002; Colgin 2016; Vinogradova 1995). MS and LS are anatomically connected with RE, ENT, RSC, (Bokor et al. 2002; Unal et al. 2015) and also participate in fear memory processing (Vetere et al. 2017; Knox et al. 2016; Tronson et al. 2009).

Despite this, we have not found significant changes in c-Fos activity in PVT, AM, AV, LD thalamic nuclei, and lateral septum of T286A^+/-^ compared to WT littermate mice in any of examined conditions (**Figure 5**).

On the other hand, we found high temporal- and valence-specific upregulation of the PVT activity, where both genotypes showed hyperexpression of c-Fos following remote extinction (**Figure 5C**), supporting its predominant role in processing fear memory at remote time-points (Do-Monte et al. 2015; Silva et al. 2018). No genotype effect, in turn, suggests that αCaMKII activity in the PVT is dispensable during extinction learning. Moreover, PVT activity is crucial for two contrasting forms of fear regulation at the same time, retrieval and extinction (Do-Monte et al. 2015; Tao et al. 2021), which can explain higher activity found in both genotypes following remote extinction. It would be then adequate to distinguish those opposite circuits within the PVT in future experiments, by labeling engram cells of fear memory and validate its activity.

Interestingly, in our paradigm, impaired extinction was associated with a prominent c-Fos increase in CM, MD, RE, AD, and MS (**Figure 5**). Such distinctive activity patterns suggest that midline, anterodorsal thalamus, and medial septum serve as hub nodes in the brain circuit of impaired remote fear extinction and their activity depends on autophosphorylation of αCaMKII-T286. Accumulating data present that RE serves as a relay for modulation of enduring, aversive memories at system levels (Wheeler et al. 2013; Vetere et al. 2017; Quet et al. 2020; Ferraris et al. 2021) through CA1-RE-mPFC pathway during recent fear extinction (Ramanathan et al. 2018) or IL-RE-BLA during remote (Silva et al. 2021). Our observations show that RE is activated when fear memory cannot be extinguished, however, similar to Silva et al. (2021) we did not see elevated RE c-Fos expression upon successful remote extinction in WT mice (**Figure 5D**). Successful contextual fear memory extinction does not activate the CM at remote time-point (Silva et al. 2018), which is supported by our observation. Nevertheless, recent cued fear memory extinction induces c-Fos in this structure (Furlong et al. 2016). The latter finding is inconsistent with our results and shows that CM is differently activated upon contextual and cued recent fear extinction. Moreover, an increase in activity of CM was noted exclusively after impaired remote fear extinction (**Figure 6A**), which suggests that CM might be involved in driving expression of fear when the animal is re-exposed to the fearful context, but this hypothesis has to be confirmed with additional experiments.

The MD is reciprocally interconnected with IL, PrL, ENT, and BLA (Tao et al. 2021; Lee et al. 2012; Zhang and Bertram 2002), and as shown by Zhang and colleagues (2002), stimulation of MD induces LTP in the BLA and ENT, which corresponds to upregulated c-Fos observed in these structures following remote extinction of T286A^+/-^ mice. This implies the possibility of the existence of PrL-MD-BLA-ENT functional connectivity upon impaired remote extinction of contextual fear. In line with this idea is the MD-BLA inactivation, which resulted in impairment of recent fear extinction. This evidence also shows that MD–BLA feedforward inhibition supports long-lasting fear attenuation (Baek et al. 2019). And finally, MD-mPFC inputs potentiation are required for the recall of remote extinction memory, supporting the notion that the engram of distant memory is scattered throughout the brain network and is possibly controlled by subcortical regions (Hugues and Garcia 2007).

Consequently, AD is one of the major inputs to RSG (van Groen and Wyss 1990; van Groen and Wyss 2003; Shibata 1993) and provides reciprocal connections to hippocampus (Wyss et al. 1979; Vetere et al. 2021). It has been shown that retrosplenial–AD theta coherence increased after recent and remote fear extinction, while gamma coherence prior to contextual fear conditioning predicted successful extinction (Corcoran et al. 2016). et al.

The recent contextual fear retrieval requires AD activity (Lopez et al. 2018), while the remote - its active suppression through long-range inhibitory projections from CA3 (Vetere et al. 2021). It is, therefore, possible that low hippocampal activity of T286A^+/-^ mice, observed here, disrupted feedback with AD and resulted in its hyperactivation upon remote extinction. As Vetere et al. (2021) reports, a similar effect was observed when inhibition of GABAergic (CA3-AD) projections resulted in upregulated c-Fos in AD only at remote time point of fear retrieval.

Impaired remote fear extinction is also accompanied by hyperactivity in the MS region, indicating a contribution of MS activity to the extinction consolidation deficits. This observation can be further supported by MS inactivation studies, showing the evidence for MS involvement in auditory (Xiao et al. 2018) and contextual fear conditioning (Calandreau et al. 2007) through adaptive processing of the contextual component or involvement in the acquisition of fear extinction (Knox and Keller 2016). Together with anatomical data, indicating dense MS projections reaching ventral and dorsal hippocampus (Amaral and Kurz 1985; Nyakas et al. 1987; Müller and Remy 2018; Khakpai et al. 2013) or ENT (Unal et al. 2015) this may suggest a putative role of MS (along with ENT and hippocampus) in processing the contextual components of the possibly dangerous environment. Another line of support is that MS-CA1 and MS-ENT connectome is involved in encoding space as shown by (Justus et al. 2017; Fuhrmann et al. 2015). However, the detailed role of the MS in remote extinction is far from clear.

Remote memory processing has been shown to engage broad hippocampal-thalamic-cortical networks (Wheeler et al. 2013; Silva et al. 2018; Vetere et al. 2021; Silva et al. 2021). Nevertheless, here we provide evidence for septal-thalamic-cortical regional co-activation upon remote fear extinction that depends on autophosphorylation of αCaMKII-T286. Such observation indicates that activation of these brain areas changes in time and might be regulated by αCaMKII activity. A similar hypothesis was proposed by Corcoran et al. (2013) who showed that PKA activation is differently engaged upon recent and remote fear extinction. Therefore, the data reported here support the notion whereby the fear extinction learning engages different neuronal networks and potentially uses different molecular mechanisms as compared to recent fear extinction. Together, our results contribute to knowledge about functioning of circuits beyond impaired fear extinction, which is of great interest both for clinical applications and for the basic understanding of memory processes. Yet, further studies including functional manipulation of septal-thalamic-cortical pathways are necessary for a more comprehensive understanding of their role in remote fear extinction.

