## Supplemental figure 1. for "Identification of neuronal ensembles involved in remote fear memory extinction impairments"

### SUPPLEMENTARY MATERIALS

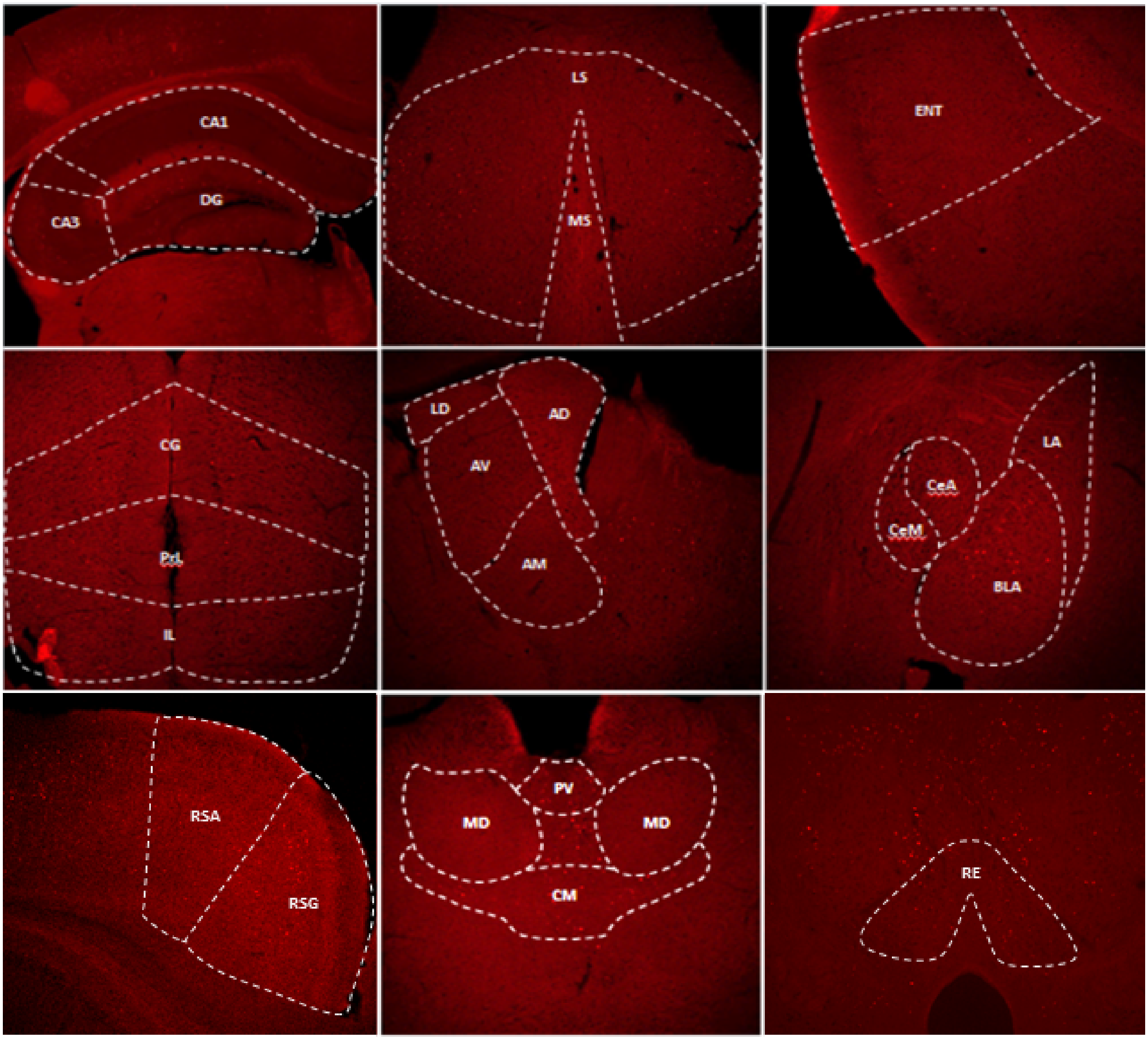


**Supplementary Figure 1.** Representative microphotographs of c-Fos$^+$ cells in all regions studied in WT animals after recent extinction.

| Brain region | WT | | | | | $T286A^{+/-}$ | | | | |
|---|---|---|---|---|---|---|---|---|---|---|
| | Naive | Ctx | 5US | Recent | Remote | Naive | Ctx | 5US | Recent | Remote |
| **dCA1** | 119.60 (36.28) | 214.40 (92.96) | 50.38 (18.18) | 35.92 (8.75) | 37.40 (19.76) | 12.61 (6.06) | 26.95 (15.60) | 58.72 (34.75) | 90.35 (60.54) | 8.61 (4.80) |
| **dCA3** | 114.46 (32.63) | 142.91 (62.41) | 64.69 (18.92) | 35.07 (11.34) | 40.39 (15.68) | 53.55 (25.56) | 37.29 (15.25) | 11.09 (4.27) | 59.96 (18.96) | 45.58 (17.06) |
| **dDG** | 63.35 (19.33) | 155.77 (24.08) | 28.61 (5.82) | 106.27 (10.74) | 90.61 (8.91) | 43.04 (12.92) | 74.24 (9.37) | 24.67 (6.09) | 67.57 (12.33) | 60.20 (12.57) |
| **BLA** | 3.45 (1.50) | 11.07 (2.53) | 2.67 (0.37) | 6.30 (1.87) | 16.44 (0.83) | 5.49 (2.17) | 8.95 (3.22) | 1.69 (0.48) | 12.11 (3.44) | 26.51 (8.07) |
| **LA** | 2.72 (0.84) | 7.59 (2.30) | 3.77 (1.38) | 7.07 (1.12) | 18.96 (3.74) | 6.01 (2.94) | 8.40 (2.26) | 4.85 (0.88) | 10.33 (2.57) | 17.29 (3.87) |
| **CeA** | 2.31 (0.71) | 5.39 (1.28) | 2.74 (0.84) | 7.29 (2.08) | 11.51 (2.89) | 5.36 (2.06) | 5.04 (0.87) | 2.04 (0.71) | 5.80 (2.58) | 8.38 (3.99) |
| **CeM** | 0.69 (0.69) | 5.58 (2.41) | 0.44 (0.44) | 7.87 (2.93) | 7.75 (2.32) | 7.2 (3.33) | 6.75 (3.01) | 0.85 (0.85) | 2.17 (1.66) | 4.91 (3.28) |
| **PrL** | 3.80 (1.65) | 30.39 (9.27) | 10.28 (3.06) | 19.31 (7.89) | 28.14 (7.03) | 12.02 (4.09) | 15.08 (4.61) | 9.14 (2.43) | 20.05 (5.00) | 58.88 (16.14) |
| **IL** | 6.25 (0.34) | 36.45 (9.50) | 7.28 (2.17) | 37.11 (8.67) | 43.24 (4.80) | 11.72 (4.87) | 34.22 (6.36) | 3.89 (1.44) | 22.37 (6.95) | 49.22 (7.21) |
| **CG** | 1.55 (1.59) | 10.43 (5.67) | 1.99 (0.71) | 8.97 (3.41) | 21.12 (4.37) | 11.45 (3.87) | 19.24 (4.73) | 6.04 (0.80) | 21.45 (3.47) | 26.03 (1.27) |
| **RSA** | 108.37 (30.86) | 83.77 (31.95) | 102.84 (33.24) | 66.31 (12.73) | 83.20 (19.53) | 30.49 (5.01) | 65.22 (18.82) | 86.02 (26.75) | 123.04 (36.03) | 120.31 (21.00) |
| **RSG** | 107.00 (20.53) | 68.47 (22.25) | 55.43 (25.39) | 61.42 (10.98) | 146.46 (35.40) | 42.35 (14.42) | 62.16 (19.26) | 61.00 (21.94) | 117.33 (31.24) | 167.90 (17.25) |
| **ENT** | 7.17 (0.88) | 25.42 (2.56) | 6.73 (1.01) | 18.89 (5.83) | 16.56 (0.55) | 9.03 (3.32) | 13.17 (2.98) | 1.80 (0.57) | 11.05 (3.90) | 30.71 (6.48) |
| **CM** | 9.61 (4.47) | 24.24 (7.91) | 5.51 (1.08) | 46.31 (15.48) | 42.25 (9.14) | 11.72 (3.25) | 41.02 (11.15) | 6.00 (3.74) | 39.29 (15.42) | 138.03 (28.34) |
| **MD** | 7.73 (3.02) | 11.87 (4.33) | 6.26 (3.26) | 18.48 (5.08) | 14.64 (1.69) | 10.56 (2.95) | 18.43 (4.74) | 7.72 (3.95) | 10.02 (3.61) | 77.17 (16.30) |
| **PV** | 26.33 (2.28) | 73.44 (30.54) | 27.40 (9.84) | 113.40 (35.02) | 264.45 (28.92) | 37.39 (20.04) | 68.06 (7.23) | 32.99 (12.64) | 111.16 (30.28) | 315.58 (91.64) |
| **RE** | 19.35 (4.76) | 45.30 (9.00) | 14.30 (4.63) | 51.30 (19.25) | 54.95 (13.70) | 29.47 (13.66) | 30.77 (8.58) | 17.23 (5.65) | 52.09 (23.56) | 112.10 (16.54) |
| **AD** | 17.74 (5.57) | 7.79 (5.21) | 8.33 (2.98) | 16.56 (4.11) | 10.74 (3.71) | 9.29 (2.89) | 4.75 (3.23) | 4.96 (2.33) | 5.15 (2.18) | 46.22 (16.64) |
| **AM** | 1.17 (1.00) | 3.98 (1.11) | 1.91 (0.95) | 1.80 (1.42) | 1.70 (1.37) | 1.19 (0.51) | 4.49 (1.11) | 2.26 (0.62) | 1.10 (0.51) | 5.04 (1.29) |
| **AV** | 2.84 (1.03) | 4.74 (1.82) | 2.47 (1.10) | 6.17 (1.49) | 6.74 (0.65) | 5.52 (1.37) | 4.32 (0.68) | 1.29 (0.64) | 3.17 (0.62) | 13.59 (4.09) |
| **LD** | 2.13 (1.31) | 0.97 (0.65) | 0.97 (0.97) | 2.93 (1.91) | 1.00 (1.00) | 1.10 (1.10) | 0.92 (0.92) | 1.53 (1.53) | 2.43 (2.43) | 7.90 (3.57) |
| **LS** | 6.69 (2.54) | 20.85 (10.91) | 4.69 (1.05) | 26.50 (6.10) | 20.83 (5.60) | 6.31 (1.71) | 28.86 (8.84) | 4.94 (1.55) | 21.57 (4.23) | 52.60 (14.87) |
| **MS** | 3.82 (1.73) | 5.73 (1.59) | 3.12 (1.62) | 4.35 (1.57) | 10.30 (3.12) | 2.97 (0.58) | 10.06 (2.57) | 2.54 (0.37) | 10.11 (3.85) | 24.23 (4.89) |

**Supplementary Table 1.** The mean value (+/-SEM) of c-Fos density (cells/mm2) captured in hippocampus (dCA1, dCA3, dDG), amygdala (BLA, LA, CeA, CeM), cortex (RSG, RSA, ENT, PrL, IL, CG), thalamus (AD, AV, AM, LD, RE, CM, MD, PV) and septal area (MS, LS) in WT mice and $T286A^{+/-}$ αCaMKII mice. Table contains results from two experimental groups: Recent, Remote extinction and three controls: Naive Ctx and 5US. CA1 - field CA1 of hippocampus, CA3 - field CA3 of hippocampus, DG - dentate gyrus, LS - lateral septum, MS - medial septum, RSG - granular part of retrosplenial cortex, RSA - agranular part of retrosplenial cortex, V1 - primary visual cortex, ENT - entorhinal cortex, PrL - prelimbic cortex, IL - infralimbic cortex, CG - cingulate cortex, AD - anterodorsal thalamic nucleus, AV - anteroventral thalamic nucleus, AM - anteromedial thalamic nucleus, LD - laterodorsal thalamic nucleus, RE - nucleus reuniens, CM - central medial thalamic nucleus, MD - mediodorsal thalamic nucleus, PV - paraventricular thalamic nucleus, BLA - anterior part of basolateral amygdaloid nucleus, LA - lateral amygdaloid nucleus, CeA - central amygdaloid nucleus, CeM - medial part of central amygdaloid nucleus

| Brain region | Genotype × Condition | Genotype | Training |
|---|---|---|---|
| **dCA1** | F(4, 39) = 2.163; (p = 0.09) | F(1, 39) = 20.62; (p < 0.001) | F(4, 39) = 2.404; (p = 0.07) |
| **dCA3** | F(4, 42) = 2.448; (p = 0.06) | F(1, 42) = 6.173; (p = 0.02) | F(4, 42) = 2.677; (p = 0.04) |
| **dDG** | F(4, 43) = 1.868; (p = 0.1) | F(1, 43) = 16.25 (p < 0.001) | F(4, 43) = 12.85; (p < 0.001) |
| **BLA** | F(4, 47) = 1.324; (p = 0.27) | F(1, 47) = 2.382; (p = 0.13) | F(4, 47) = 11.26; (p < 0.001) |
| **LA** | F(4, 47) = 0.377; (p = 0.82) | F(1, 47) = 0.874; (p = 0.35) | F(4, 47) = 11.13; (p < 0.001) |
| **CeA** | F(4, 46) = 1.318; (p = 0.28) | F(1, 46) = 0.496; (p = 0.49) | F(4, 46) = 5.452; (p = 0.001) |
| **CeM** | F(4, 44) = 1.778; (p = 0.151) | F(1, 44) = 0.004; (p = 0.953) | F(4, 44) = 2.15; (p = 0.091) |
| **PrL** | F(4, 42) = 2.659; (p = 0.05) | F(1, 42) = 1.374; (p = 0.42) | F(4, 42) = 7.534; (p < 0.001) |
| **IL** | F(4, 40) = 1.02; (p = 0.41) | F(1, 40) = 0.074; (p = 0.79) | F(4, 40) = 15.09; (p < 0.001) |
| **CG** | F(4, 40) = 1.308; (p = 0.28) | F(1, 40) = 0.522; (p = 0.47) | F(4, 40) = 10.84; (p < 0.001) |
| **RSA** | F(4, 43) = 2.199; (p = 0.085) | F(1, 43) = 0.058; (p = 0.811) | F(4, 43) = 0.609; (p = 0.658) |
| **RSG** | F(4, 42) = 1.956; (p = 0.12) | F(1, 42) = 0.027; (p = 0.87) | F(4, 42) = 5.494; (p = 0.001) |
| **ENT** | F(4, 41) = 4.168; (p = 0.006) | F(1, 41) = 5.317; (p = 0.026) | F(4, 41) = 13.46; (p < 0.001) |
| **CM** | F(4, 44) = 6.138; (p = 0.001) | F(1, 44) = 9.129; (p = 0.004) | F(4, 44) = 16.13; (p < 0.001) |
| **MD** | F(4, 44) = 12.28; (p < 0.001) | F(1, 44) = 14.79; (p < 0.001) | F(4, 44) = 15.48; (p < 0.001) |
| **PVT** | F(4, 44) = 0.249; (p = 0.91) | F(1, 44) = 0.388; (p = 0.54) | F(4, 44) = 22.08; (p < 0.001) |
| **RE** | F(4, 40) = 3.452; (p = 0.02) | F(1, 40) = 2.971; (p = 0.09) | F(4, 40) = 7.344; (p < 0.001) |
| **AD** | F(4, 39) = 3.54; (p = 0.01) | F(1, 39) = 0.144; (p = 0.71) | F(4, 39) = 2.999; (p = 0.03) |
| **AM** | F(4, 37) = 1.647; (p = 0.18) | F(1, 37) = 1.701; (p = 0.2) | F(4, 37) = 3.262; (p = 0.022) |
| **AV** | F(4, 39) = 1.867; (p = 0.14) | F(1, 39) = 0.575; (p = 0.45) | F(4, 39) = 4.443; (p = 0.005) |
| **LD** | F(4, 35) = 0.597; (p = 0.67) | F(1, 35) = 1.059; (p = 0.31) | F(4, 35) = 8.112; (p < 0.001) |
| **LS** | F(4, 39) = 1.845; (p = 0.14) | F(1, 39) = 2.043; (p = 0.16) | F(4, 39) = 6.183; (p = 0.001) |
| **MS** | F(4, 35) = 2.409; (p = 0.07) | F(1, 35) = 6.214; (p = 0.018) | F(4, 35) = 8.927; (p < 0.001) |

**Supplementary Table 2.** The main two-way ANOVA F-test results for each analysed brain structure.
